# An early cortical progenitor-specific mechanism regulates thalamocortical innervation

**DOI:** 10.1101/2020.09.25.312785

**Authors:** Suranjana Pal, Deepanjali Dwivedi, Tuli Pramanik, Geeta Godbole, Takuji Iwasato, Denis Jabaudon, Upinder S. Bhalla, Shubha Tole

## Abstract

The cortical subplate is critical in regulating the entry of thalamocortical sensory afferents into the cortex. These afferents reach the subplate at embryonic day (E)15.5 in the mouse, but “wait” for several days, entering the cortical plate postnatally. We report that when transcription factor Lhx2 is lost in E11.5 cortical progenitors, which give rise to subplate neurons, thalamocortical afferents display premature, exuberant innervation of the E15.5 cortex. Embryonic mutant subplate neurons are correctly positioned below the cortical plate, but they display an altered transcriptome and immature electrophysiological properties during the waiting period. The sensory thalamus in these cortex-specific *Lhx2* mutants displays atrophy, eventually leading to severe deficits in thalamocortical innervation. Strikingly, these phenotypes do not manifest if Lhx2 is lost in postmitotic subplate neurons. These results demonstrate a mechanism operating in subplate progenitors that has profound consequences on the growth of thalamocortical axons into the cortex.

## Introduction

Sensory input reaches the neocortex via thalamocortical tract, and the precise topography and connectivity of this tract is critical to sensory processing. A key feature of cortical innervation by thalamocortical axons is a “waiting period” during which thalamic afferents enter the dorsal telencephalon while cortical neurogenesis is still underway, but “wait” within the cortical subplate for a period lasting several days in rodents, before innervating layer 4 of the cortex (1–6). The importance of the subplate as an intermediate target of the thalamocortical afferents is well established from studies in which subplate neurons were either experimentally ablated, or mispositioned/reduced due to genetic perturbations [reviewed in (7)]. Thalamocortical afferents make synaptic contacts with the subplate (5), and electrical activity in the subplate is critical for correct thalamocortical innervation of the cortical plate at later stages (8–11). Subplate neurons are born at embryonic day (E) 11.5 in the mouse from common progenitors that will later produce the neurons of the cortical plate (12). However, the mechanisms that regulate the properties of the subplate neurons that are necessary for the waiting period are unclear, because the regulatory processes in cortical progenitors that give rise to subplate neurons are poorly understood.

Previously, we reported a loss of the somatosensory barrels upon cortex-specific deletion of *Lhx2* (13). Barrels are a prominent feature of the rodent somatosensory cortex, formed by local aggregates of layer 4 cortical neurons when thalamocortical afferents innervate the cortex (14). We found that the barrels are absent when *Lhx2* is disrupted using an Emx1Cre line which acts in cortical progenitors from E11.5 [Emx1Cre^11.5^; (13)]. In contrast, loss of *Lhx2* using the postmitotic neuron-specific NexCre permits somatosensory barrels to be formed in layer 4 (15,16). Based on these two distinct phenotypes, we hypothesized that Lhx2 function in cortical progenitors is required for the key aspects of thalamocortical axon development.

Here, we analyze thalamocortical axon projection defects when *Lhx2* is disrupted in cortical progenitors using Emx1Cre^11.5^. We find that, although there is extremely limited thalamocortical innervation detectable in the postnatal cortex of *Emx1Cre*^*11*.*5*^;*Lhx2*^*lox/lox*^ mice, there is a premature exuberant innervation of the cortical plate in the embryo at E15.5, such that thalamocortical fibers project beyond the subplate and reach the marginal zone. This is followed by atrophy of the prominent sensory ventrobasal nucleus (VB) of the thalamus, and eventual loss of thalamocortical innervation of the cortex. Using CreER-driven tamoxifen-inducible *Lhx2* deletion, we identify that loss of Lhx2 from (E11.5), but not from E12.5 or E13.5, recapitulates the aberrant innervation phenotype, implicating the subplate in this process, since it is born at E11.5. The subplate in *Emx1Cre*^*11*.*5*^;*Lhx2*^*lox/lox*^ brains expresses specific markers and occupies its appropriate position under the cortical plate but exhibits severe dysregulation of its transcriptome. In addition, the electrophysiological properties of these subplate neurons are perturbed compared with controls. Our results offer novel insights into mechanisms that operate in cortical progenitors that are necessary for proper subplate differentiation and for thalamocortical axon development and circuit assembly.

## Results

In order to visualize thalamocortical projections, we used a TCA-GFP reporter line in which the expression of membrane-bound enhanced GFP is driven in primary sensory thalamic nuclei under serotonin transporter (*SERT*) promoter (17). The TCA-GFP background allowed us to compare the two *Lhx2* conditional mutants that were previously reported to have defects in thalamocortical innervation: a complete loss of somatosensory barrels [*Emx1Cre*^*11*.*5*^;*Lhx2*^*lox/lox*^; (13)]; or defects in polarized dendritic extension of layer 4 spiny stellate neurons [*NexCre;Lhx2*^*lox/lox*^; (16)].

Postnatal day (P) 7 brains were examined for GFP fluorescence indicative of thalamocortical innervation in either tangential slices of the cortex or coronal sections of hemispheres. The barrel map was visible in control brains (Figure 1A), but was profoundly disrupted in *Emx1Cre*^*11*.*5*^;*Lhx2*^*lox/lox*^ brains, in which tangential slices revealed extremely reduced innervation in unstructured clusters that did not form any recognizable barrel field pattern (arrowheads, Figure 1B). In contrast, barrels were still apparent in *NexCre;Lhx2*^*lox/lox*^ brains (Figure 1C), confirming that postmitotic Lhx2 is not required for barrel patterning. Coronal sections of *Emx1Cre*^*11*.*5*^;*Lhx2*^*lox/lox*^ brains showed the GFP-expressing axons to be clustered at one location and projecting aberrantly up to the marginal zone of the cortex, instead of layer 4. Quantification of the fluorescence intensity in the somatosensory cortex revealed a significant reduction in *Emx1Cre*^*11*.*5*^;*Lhx2*^*lox/lox*^ brains compared with controls (Figure 1D).

**Figure 1:**
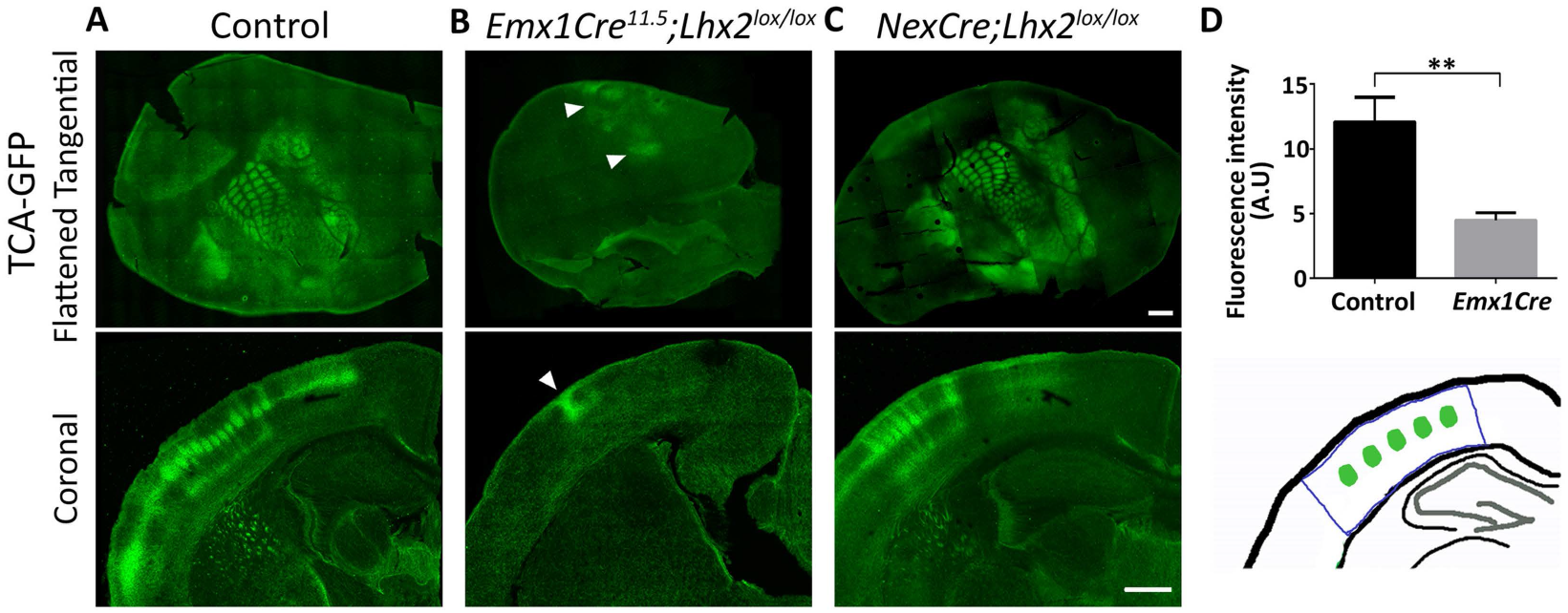
Loss of Lhx2 mediated by Emx1Cre causes severe disruption of somatosensory barrels. (A) TCA-GFP reporter expression reveals the thalamocortical innervation and somatosensory barrels in flattened tangential sections and coronal sections at P7 in control brains. (B) In *Emx1Cre*^*11*.*5*^;*Lhx2*^*lox/lox*^;*TCA-GFP* brains, aberrant thalamocortical innervation (arrowheads) is seen clustered in a limited region of the cortex at P7. (C) *NexCre;Lhx2*^*lox/lox*^;*TCA-GFP* brains reveal that postmitotic deletion of *Lhx2* permits thalamocortical afferents to innervate layer 4 and barrels can be detected. (D) Fluorescence intensity of the somatosensory cortex, measured in coronal sections, is significantly reduced in P7 *Emx1Cre*^*11*.*5*^;*Lhx2*^*lox/lox*^ brains compared with controls. n= 3 brains per condition. Tangential and coronal TCA-GFP images in A-C are composites of multiple confocal images. All scale bars are 500 µm.

The VB nucleus of the thalamus projects to the barrel cortex, and its neurons require the establishment of stable synapses with their layer 4 targets for survival (18,19). Therefore, we examined whether atrophy of the thalamic nuclei may explain the impoverished thalamocortical projections in the P7 cortex of *Emx1Cre*^*11*.*5*^;*Lhx2*^*lox/lox*^ mutant brains. Indeed, the VB was greatly shrunken in mutant brains, as seen by the expression of the serotonin transporter [*SERT*; Figure 2A-C; (20)]. Interestingly, this VB shrinkage was apparent from E17.5, when thalamocortical axons are still in their waiting period (Figure 2A-C) and corticothalamic projections from layer 6 have yet to enter the VB (Figure 2D; (21,22). Together, these results suggest that the VB may require a cortex-derived, Lhx2-regulated signal which is available to thalamocortical axons during the waiting period. The shrinkage of the VB provides an explanation for the postnatal phenotype in *Emx1Cre*^*11*.*5*^;*Lhx2*^*lox/lox*^ mutant brains [Figure 1; (13)].

**Figure 2:**
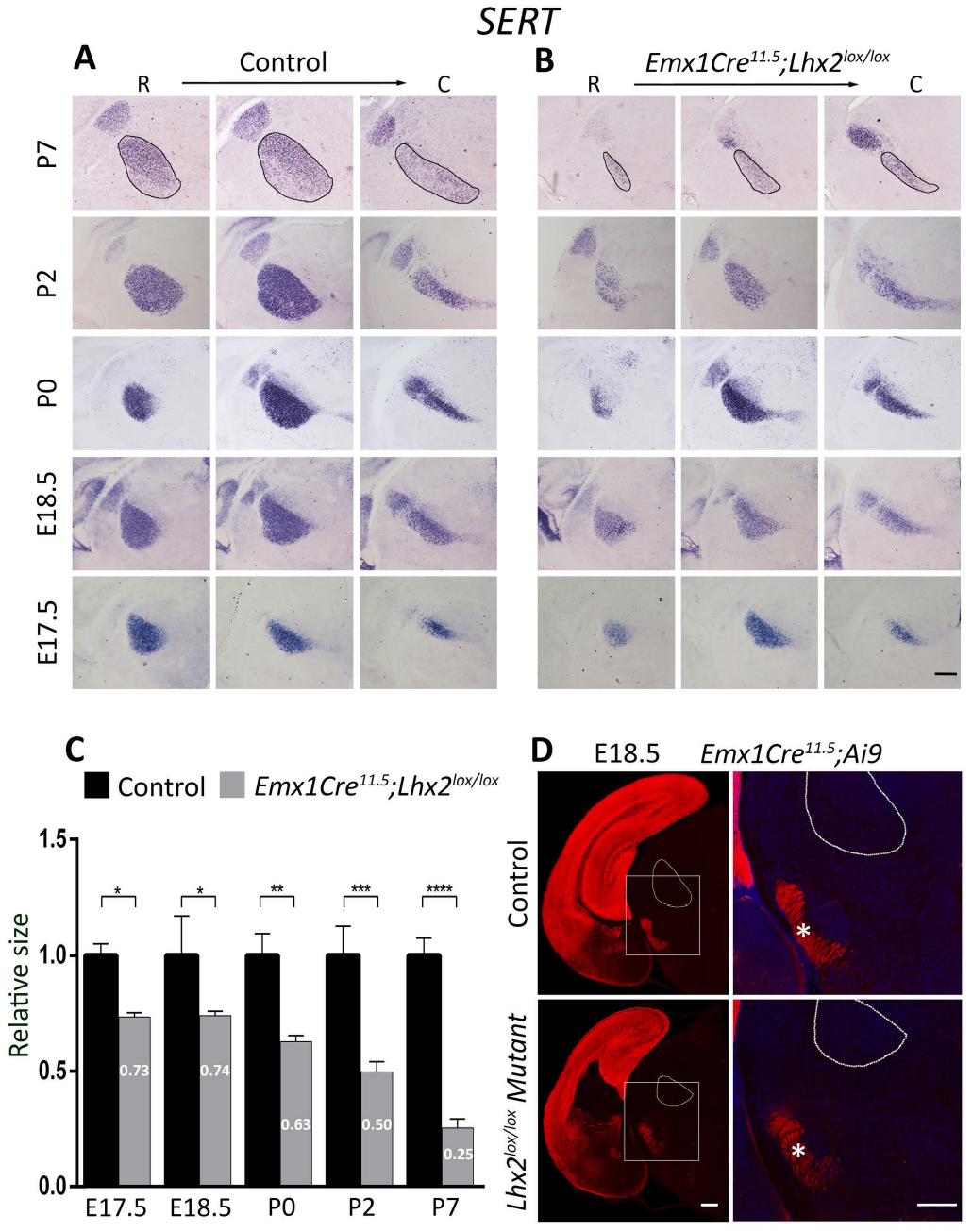
Loss of Lhx2 mediated by Emx1Cre causes shrinkage of the somatosensory VB nucleus from embryonic stages. (A, B) Expression of *SERT* in the thalamus of control and *Emx1Cre*^*11*.*5*^;*Lhx2*^*lox/lox*^ brains through embryonic and postnatal development (E17.5-P7). A rostro-caudal series is shown for each stage. The somatosensory VB nucleus (black ovals) appears smaller in *Emx1Cre*^*11*.*5*^;*Lhx2*^*lox/lox*^ brains compared with controls at each stage examined. (C) Quantification of VB area. A comparison of the average values from control and *Emx1Cre*^*11*.*5*^;*Lhx2*^*lox/lox*^ brains reveals the VB to be significantly smaller in *Emx1Cre*^*11*.*5*^;*Lhx2*^*lox/lox*^ brains at each age examined. n=3 brains per condition. (D) An Ai9 reporter reveals the corticothalamic axons in control and *Emx1Cre*^*11*.*5*^;*Lhx2*^*lox/lox*^ brains have reached the reticular thalamic nucleus at E18.5 (white asterisks), but have not yet innervated the VB nucleus (white ovals). The boxed region is shown at high magnification together with the DAPI channel (blue). All scale bars are 500 µm.

The finding that the VB appears to be affected as early as E17.5 in *Emx1Cre*^*11*.*5*^;*Lhx2*^*lox/lox*^ mutants motivated the hypothesis that thalamocortical axons may display deficits in their development during the waiting period. Since the TCA-GFP line does not display robust fluorescence at embryonic stages, we used *in utero* electroporation to label embryonic VB projections by transfecting a GFP-expressing plasmid into the diencephalon of control and mutant embryos at E11.5, when VB neurons are born. Control, *Emx1Cre*^*11*.*5*^;*Lhx2*^*lox/lox*^, and *NexCre;Lhx2*^*lox/lox*^ brains were examined at E15.5, by which stage the GFP-expressing thalamocortical tract has entered the dorsal telencephalon (Figure 3A). Cortical thickness was similar in control, *Emx1Cre*^*11*.*5*^;*Lhx2*^*lox/lox*^, and *NexCre;Lhx2*^*lox/lox*^ brains (Figure 3B). In contrast to control embryos, in which the thalamocortical tract courses below the cortical plate in a tight bundle, *Emx1Cre*^*11*.*5*^;*Lhx2*^*lox/lox*^ brains displayed an exuberant outgrowth of thalamocortical fibers into the cortical plate at E15.5 (Figure 3C, D). Furthermore, thalamocortical axons appeared defasciculated, extended several branches prematurely into the cortical plate, and extended aberrantly upto the marginal zone in *Emx1Cre*^*11*.*5*^;*Lhx2*^*lox/lox*^ brains (arrowheads, Figure 3C). Such exuberant branching was not seen in E15.5 *NexCre;Lhx2*^*lox/lox*^ brains, which displayed a well-ordered, bundled thalamocortical tract that coursed below the cortical plate, similar to controls (Figure 3E, F). The area of the cortical plate in which green fibers were present was scored in each condition and found to be significantly increased in E15.5 *Emx1Cre*^*11*.*5*^;*Lhx2*^*lox/lox*^ brains compared with *NexCre;Lhx2*^*lox/lox*^ and control brains (Figure 3G).

**Figure 3:**
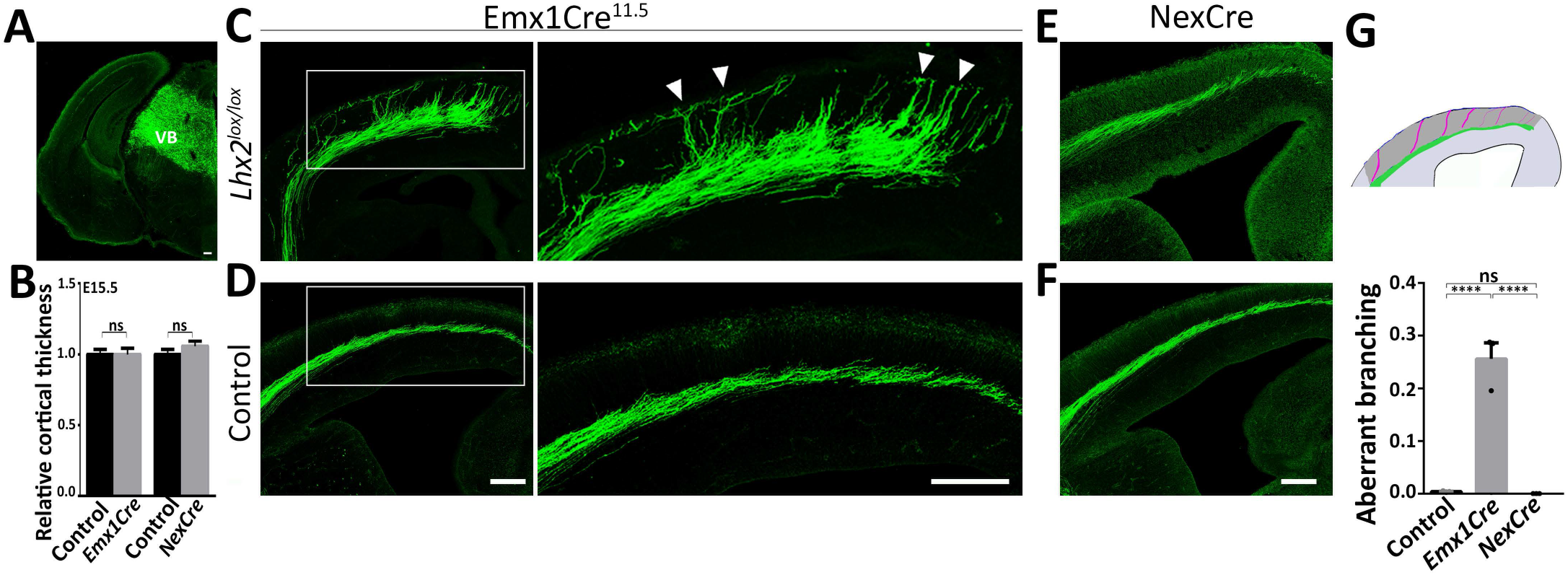
Embryonic thalamocortical axons are defasciculated and display premature growth into the cortical plate when *Lhx2* is disrupted using Emx1Cre, but not NexCre. (A) Electroporation of a plasmid encoding GFP into the diencephalon at E11.5 labels the thalamus and also thalamocortical axons by E15.5. (B) Cortical thickness is unaffected at E15.5 in *Emx1Cre*^*11*.*5*^;*Lhx2*^*lox/lox*^ and *NexCre;Lhx2*^*lox/lox*^ brains. 3 brains were analyzed per condition. The average cortical thickness from the mutant sections was normalized to that of the respective controls. (C, D) Loss of *Lhx2* using Emx1Cre causes thalamocortical afferents to defasciculate as they course through the dorsal telencephalon at E15.5, whereas they remain tightly fasciculated in controls. In *Emx1Cre*^*11*.*5*^;*Lhx2*^*lox/lox*^ brains, thalamocortical axons enter the cortical plate prematurely and extend aberrantly up to the marginal zone (arrowheads). (E, F) When *Lhx2* is disrupted using NexCre, thalamocortical afferents appear indistinguishable from those in the controls in E15.5 brains. (G) Sections of *Emx1Cre*^*11*.*5*^;*Lhx2*^*lox/lox*,^ *NexCre;Lhx2*^*lox/lox*^ and control E15.5 brains were scored for aberrant axonal branching. Scatter dots in the bar plot represent aberrant branching value for each brain (n=3 for each condition). All fluorescence images are composites of multiple confocal images. Boxes in C and D represent regions displayed at high magnification in the adjacent panels. All scale bars are 100 µm.

The thalamocortical exuberant innervation phenotype was only seen when *Lhx2* is disrupted using Emx1Cre. Therefore, we reasoned that the critical requirement for this transcription factor must be in progenitors, not postmitotic neurons. Since essentially all cortical layers are produced sequentially from progenitors that express Emx1Cre, we sought to narrow down the stage at which Lhx2 function is relevant for the proper guidance of thalamocortical axons, using tamoxifen-inducible CreERT2. Embryos of the genotype *CreERT2;Lhx2*^*lox/lox*^ were electroporated with a GFP-expressing plasmid at E11.5 in the diencephalon to label the VB thalamocortical fibers, tamoxifen was administered at different stages to disrupt *Lhx2* (Figure 4C-F), and the embryos were harvested at E15.5. Whereas tamoxifen administration at E11.5 closely recapitulated the defasciculation and exuberant innervation of thalamocortical axons seen in *Emx1Cre*^*11*.*5*^;*Lhx2*^*lox/lox*^ embryos (Figure 4B, C, G), tamoxifen administration at E12.75 and at E13.5 resulted in a phenotype similar to control brains (Figure 4A, E, F). Tamoxifen administration at E12.5 produced a phenotype mostly resembling that of control brains, except for some fibers at dorsomedial levels that reached the marginal zone of the cortex (Figure 4D). The neurons that populate the subplate are born in a latero-medial gradient, from E11.5 to E12.5, with the medial subplate being born last (23,24). Therefore, the phenotype seen upon tamoxifen administration at E12.5 to *CreERT2;Lhx2*^*lox/lox*^ embryos is consistent with the birth gradient of subplate neurons.

**Figure 4:**
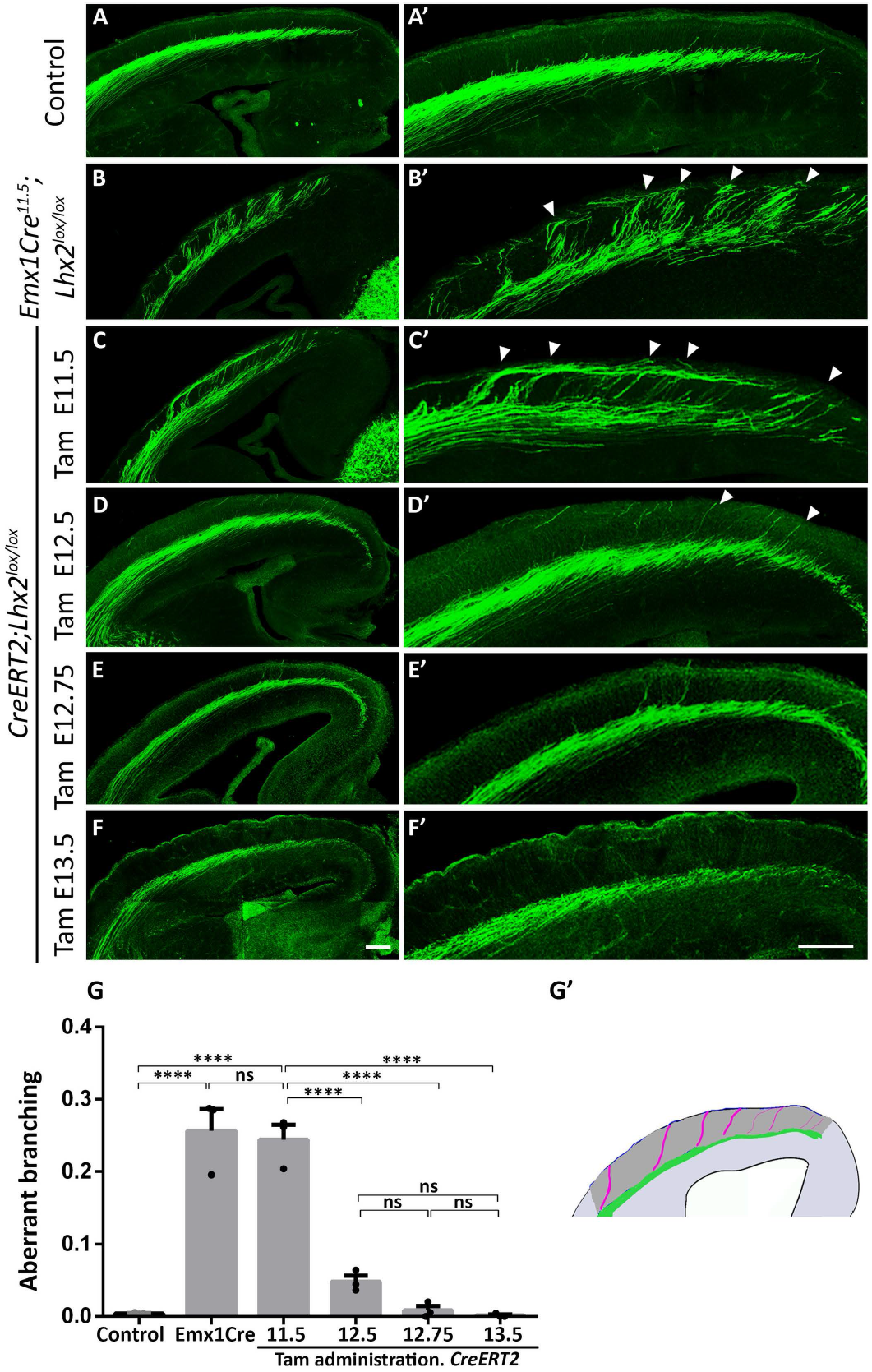
Early, but not late cortical progenitors require Lhx2 for normal thalamocortical innervation. (A-F; A’-F’) Sections of E15.5 Control (A, A’), *Emx1Cre*^*11*.*5*^;*Lhx2*^*lox/lox*^ (B, B’), *CreERT2;Lhx2*^*lox/lox*^ (C-F; C’-F’) brains in which thalamocortical innervation is visualized by electroporation of a plasmid encoding GFP into the thalamus at E11.5 (n=3 electroporated brains per condition). (A, A’) Thalamocortical afferents in control brains course in a tight bundle below the cortical plate (CP). (B, B’) Loss of *Lhx2* using Emx1Cre causes premature innervation of thalamocortical afferents that extend up to the marginal zone (arrowheads). (C-F; C’-F’) This phenotype is recapitulated when tamoxifen is administered at E11.5 to *CreERT2;Lhx2*^*lox/lox*^ but not E12.5, E12.75 or E13.5 (D-F). (G, G’) Aberrant axonal branching of TCAs seen in *CreERT2;Lhx2*^*lox/lox*^ Tam 11.5 brains is similar to *Emx1Cre*^*11*.*5*^;*Lhx2*^*lox/lox*^ brains. The exuberant branching phenotype is significantly reduced when tamoxifen is administered at E12.5, E12.75 or E13.5. Scatter dots in the bar plot represent aberrant branching value for each brain (n=3 brains per condition). All panels are composites of multiple confocal images. A’-F’ are high magnification images of A-F respectively. All scale bars are 100 µm.

In summary, these results demonstrate that Lhx2 is necessary in early (E11.5) progenitors at the time the subplate is being generated in order for normal thalamocortical axon projection to occur. This suggests that daughter subplate neurons may develop aberrantly when Lhx2 is absent in their mother cells, leading to abnormal subplate properties.

The phenotype of premature thalamocortical innervation and fibers reaching the marginal zone in the *Emx1Cre*^*11*.*5*^;*Lhx2*^*lox/lox*^ mutant was reminiscent of the thalamocortical projection phenotype described in *reeler* mutants, in which the subplate is mislocalized above the cortical plate (25). Therefore, we examined whether the cortical subplate was specified and present in its correct location in mutant brains. We first examined the expression of *CTGF* (Connective Tissue Growth Factor) and Cplx3 (Complexin 3), established markers of the postnatal subplate (26). At P7, control brains displayed a tightly packed subplate expressing both markers (Figure 5A). In *Emx1Cre*^*11*.*5*^;*Lhx2*^*lox/lox*^ brains, the subplate also expressed both markers (Figure 5B). Furthermore, subplate cells were positioned normally, below the cortical plate, though they were not as tightly packed as in the controls. Neither *CTGF* nor Cplx3 are expressed in the embryonic subplate, so we used *Nurr1* (Nr4a2; Nuclear Receptor Subfamily 4 Group A Member 2), which labels the subplate at E15.5 (Figure 5C, D). *Nurr1* was expressed in the *Emx1Cre*^*11*.*5*^;*Lhx2*^*lox/lox*^ subplate, and shows that it is correctly positioned below the cortical plate when the thalamocortical axons arrive. Therefore, the mutant phenotype of premature innervation of the cortical plate, with axons extending up to the marginal zone, could not be explained by a mislocalized subplate. We examined serial sections for *Lhx2* expression using a probe designed specifically against the floxed exon and confirmed that *Lhx2* expression was indeed lost from the entire cortex as expected for Emx1Cre-driven recombination (Figure 5 C, D). In summary, the aberrant thalamocortical innervation phenotype could not be explained by a mispositioning of subplate neurons in *Emx1Cre*^*11*.*5*^;*Lhx2*^*lox/lox*^ brains.

**Figure 5:**
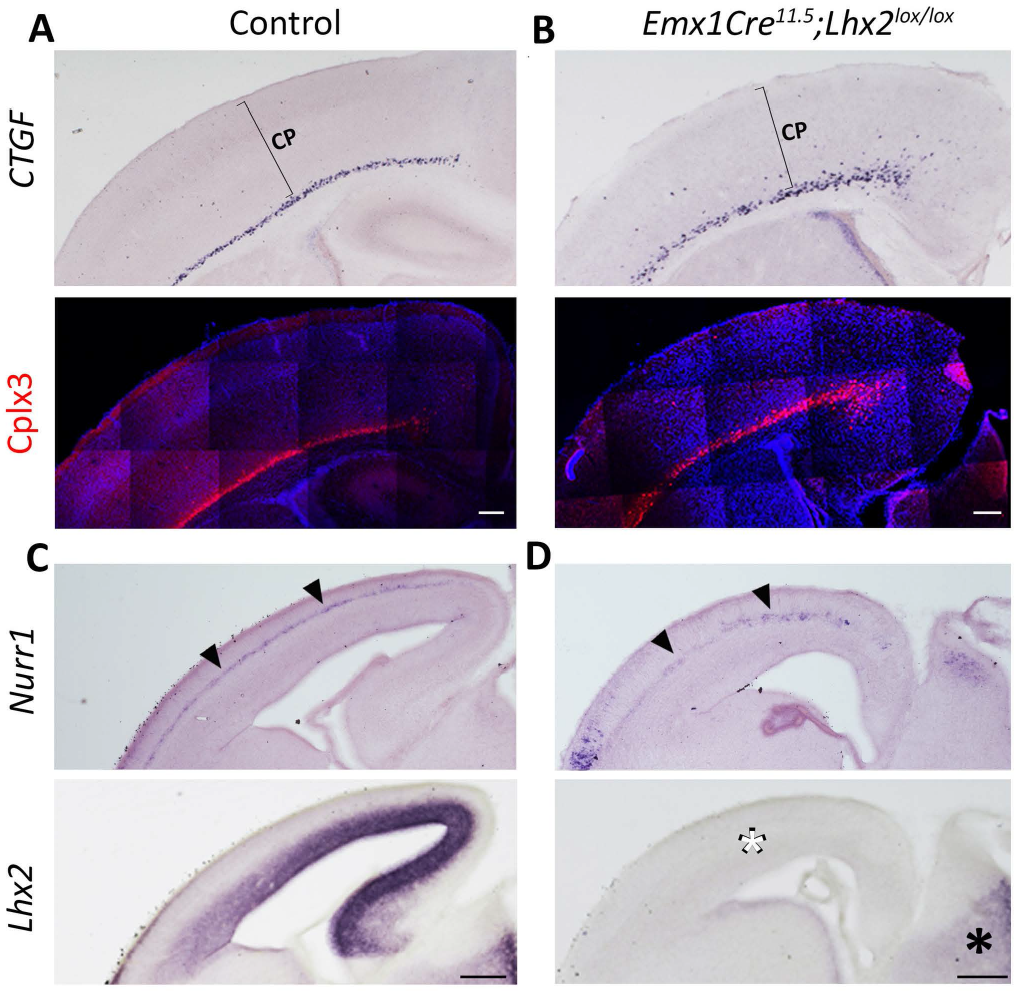
Subplate neurons are specified and positioned below the cortical plate upon Emx1Cre-mediated loss of Lhx2. (A, B) Expression of subplate markers in control and *Emx1Cre*^*11*.*5*^;*Lhx2*^*lox/lox*^ brains (n=3 brains per condition). At P7, *CTGF* expression and Cplx3 immunoreactivity reveal a tightly packed subplate in controls and a less compact subplate in mutant brains, located below the cortical plate (CP). (C, D) At E15.5, *Nurr1* is expressed in both control and mutant subplate neurons (arrowheads). *Lhx2* expression is lost in the cortex of *Emx1Cre*^*11*.*5*^;*Lhx2*^*lox/lox*^ brains (white asterisk), while expression in the thalamus is maintained (black asterisk). The Cplx3 immunohistochemistry panels in A and B are composites of multiple confocal images. All scale bars are 200 µm.

Although the position of the subplate in *Emx1Cre*^*11*.*5*^;*Lhx2*^*lox/lox*^ brains is not altered, it may be molecularly disrupted in a manner that is critical for thalamocortical axon guidance. To get an overall picture of the transcriptomic changes within subplate neurons upon loss of Lhx2, we performed RNAseq on microdissected subplate tissue. We hypothesized that loss of Lhx2 from the progenitor stage versus the postmitotic stage may have distinct regulatory consequences on the transcriptome of subplate neurons, and these differences may underlie the differences in the thalamocortical axon phenotype in *Emx1Cre*^*11*.*5*^;*Lhx2*^*lox/lox*^ and *NexCre;Lhx2*^*lox/lox*^ brains. Therefore, we compared the wild-type subplate transcriptome with that of the *Emx1Cre*^*11*.*5*^;*Lhx2*^*lox/lox*^ and *NexCre;Lhx2*^*lox/lox*^ subplate. We selected P0 as the stage of analysis because the last wave of cortical neurons has migrated past the subplate by this stage and would not be present in the dissected sample (Figure 6A). As expected, the transcriptomes of the *Emx1Cre*^*11*.*5*^;*Lhx2*^*lox/lox*^ and *NexCre;Lhx2*^*lox/lox*^ subplate were each dysregulated compared with that of their corresponding littermate controls. We examined the genes that displayed a fold change greater than 1.5 in each condition and found that 357 genes were differentially expressed in the *Emx1Cre*^*11*.*5*^;*Lhx2*^*lox/lox*^ subplate and 196 in the *NexCreLhx2*^*lox/lox*^ subplate (Figure 6B). However, when we compared these two differentially expressed datasets, only 29 genes were in common between the two (Figure 6B). This indicated that though both Cre drivers resulted in a loss of Lhx2 from the subplate, using a driver that acts in progenitors versus postmitotic cells has vastly different consequences on the transcriptome of the resulting subplate neurons.

**Figure 6:**
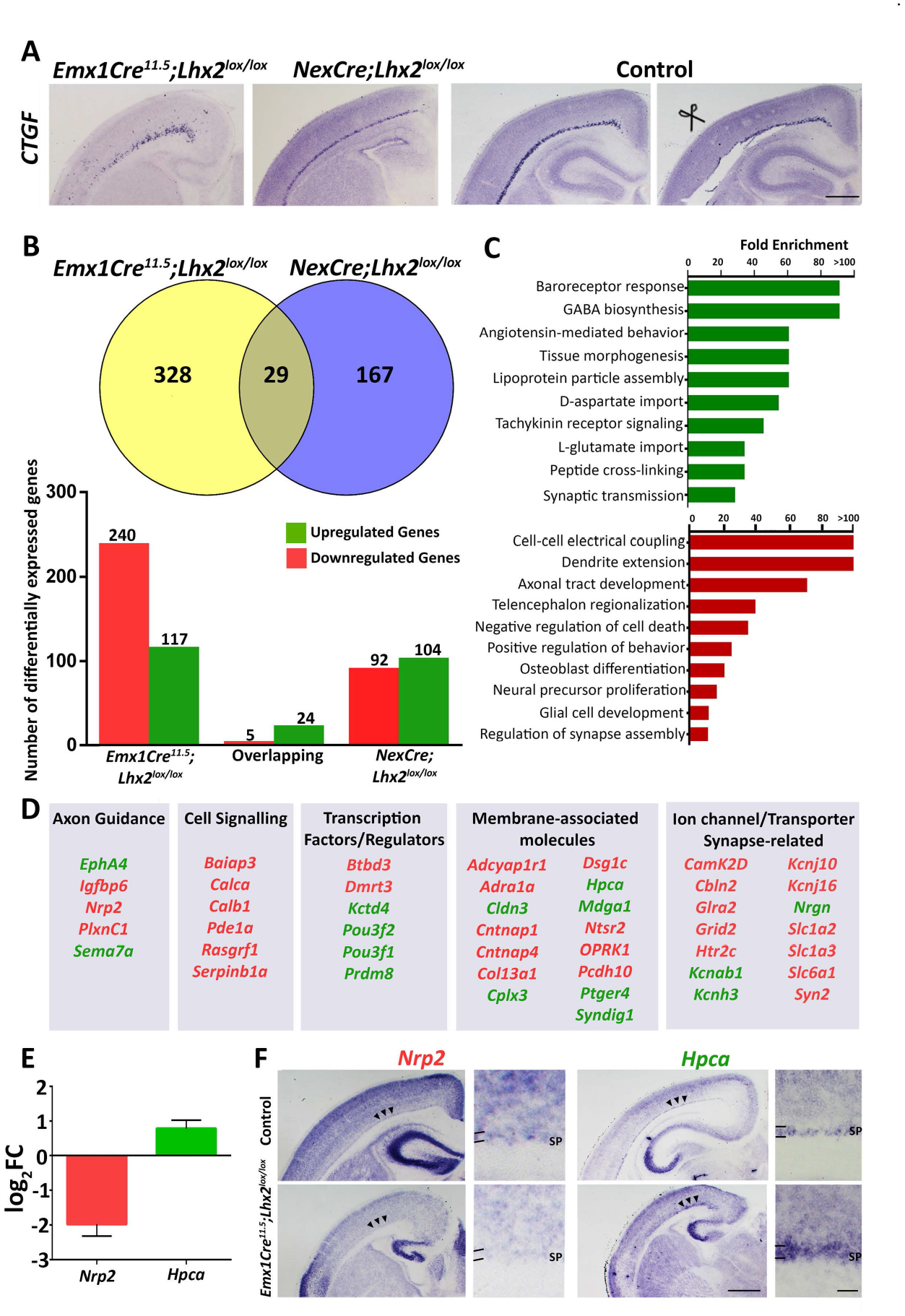
Transcriptomic analysis of the subplate. (A) *Emx1Cre*^*11*.*5*^;*Lhx2*^*lox/lox*^, *NexCre;Lhx2*^*lox/lox*^ and control sections at P0 displaying *CTGF* expression in the subplate, and a control section in which the subplate was removed by microdissection. (B) Bulk RNA seq revealed 328 and 167 differentially expressed genes unique to the *Emx1Cre*^*11*.*5*^;*Lhx2*^*lox/lox*^, *NexCre;Lhx2*^*lox/lox*^ subplate respectively, and 29 overlapping differentially expressed genes. (C, D) GO analysis of the differentially expressed genes in the *Emx1Cre*^*11*.*5*^;*Lhx2*^*lox/lox*^ subplate (C) and selected genes in different functional categories (D; See Table 1-2 and Supplementary Figure 2 for related data). (E) Bar plot representing log_2_fold change values of representative differentially expressed genes *Nrp2* and *Hpca*. (F) *Nrp2* expression is decreased and *Hpca* expression is increased in the *Emx1Cre*^*11*.*5*^;*Lhx2*^*lox/lox*^ subplate (arrowheads). SP, subplate. Scale bars in A, F are 500 µm and 50 µm for high magnification images in F.

We further analyzed the 328 differentially expressed genes that were specific to the *Emx1Cre*^*11*.*5*^;*Lhx2*^*lox/lox*^ dataset. GO analysis revealed multiple developmentally significant gene clusters (Figure 6C). We categorized these 328 genes according to their known functions and found up- and down-regulated genes that mediate axon guidance, cell signaling, transcription factors, membrane-associated molecules, as well as ion channels/transporters/synaptic molecules (Figure 6D; Supplementary Figure 2). Further, we used *in situ* hybridization to validate the RNA-seq results for some of these genes (Figure 6E, F).

**Table 1:** Gene Ontology Analysis of Differentially Regulated Genes.

| **Upregulated Genes** | **Fold Enrichment** | **p value** |
| --- | --- | --- |
| Baroreceptor response (GO:0001983) | 91.25 | 0.00171 |
| GABA biosynthesis (GO:0009449) | 91.25 | 0.0017 |
| Angiotensin-mediated behavior (GO:0003051) | 60.83 | 0.0375 |
| Tissue morphogenesis (GO:0060414) | 60.83 | 0.0374 |
| Lipoprotein particle assembly (GO:0090107) | 60.83 | 0.0373 |
| D-aspartate import (GO:0070779) | 54.75 | 0.0385 |
| Tachykinin receptor signaling (GO:0007217) | 45.63 | 0.00529 |
| L-glutamate import (GO:0098712) | 34.22 | 0.00938 |
| Peptide cross-linking (GO:0019800) | 34.22 | 0.00935 |
| Synaptic transmission (GO:0032230) | 28.52 | 0.00028 |

| **Downregulated Genes** | **Fold Enrichment** | **p value** |
| --- | --- | --- |
| Cell-cell electrical coupling (GO:0086064) | >100 | 0.0492 |
| Dendrite extension (GO:1903860) | >100 | 0.0487 |
| Axonal tract development (GO:0021957) | 72.06 | 0.00571 |
| Telencephalon Regionalization (GO:0021978) | 40.53, | 0.0205 |
| Negative regulation of cell death (GO:1903206) | 25.94 | 0.047 |
| Positive regulation of behavior (GO:0048520) | 21.62 | 0.0129 |
| Osteoblast differentiation (GO:0045668) | 16.95 | 0.0248 |
| Neural precursor proliferation (GO:2000177) | 11.48 | 0.00539 |
| Glial cell development (GO:0021782) | 9.56 | 0.0386 |
| Regulation of synapse assembly (GO:0051963) | 9.01 | 0.0478 |

**Table 2:**
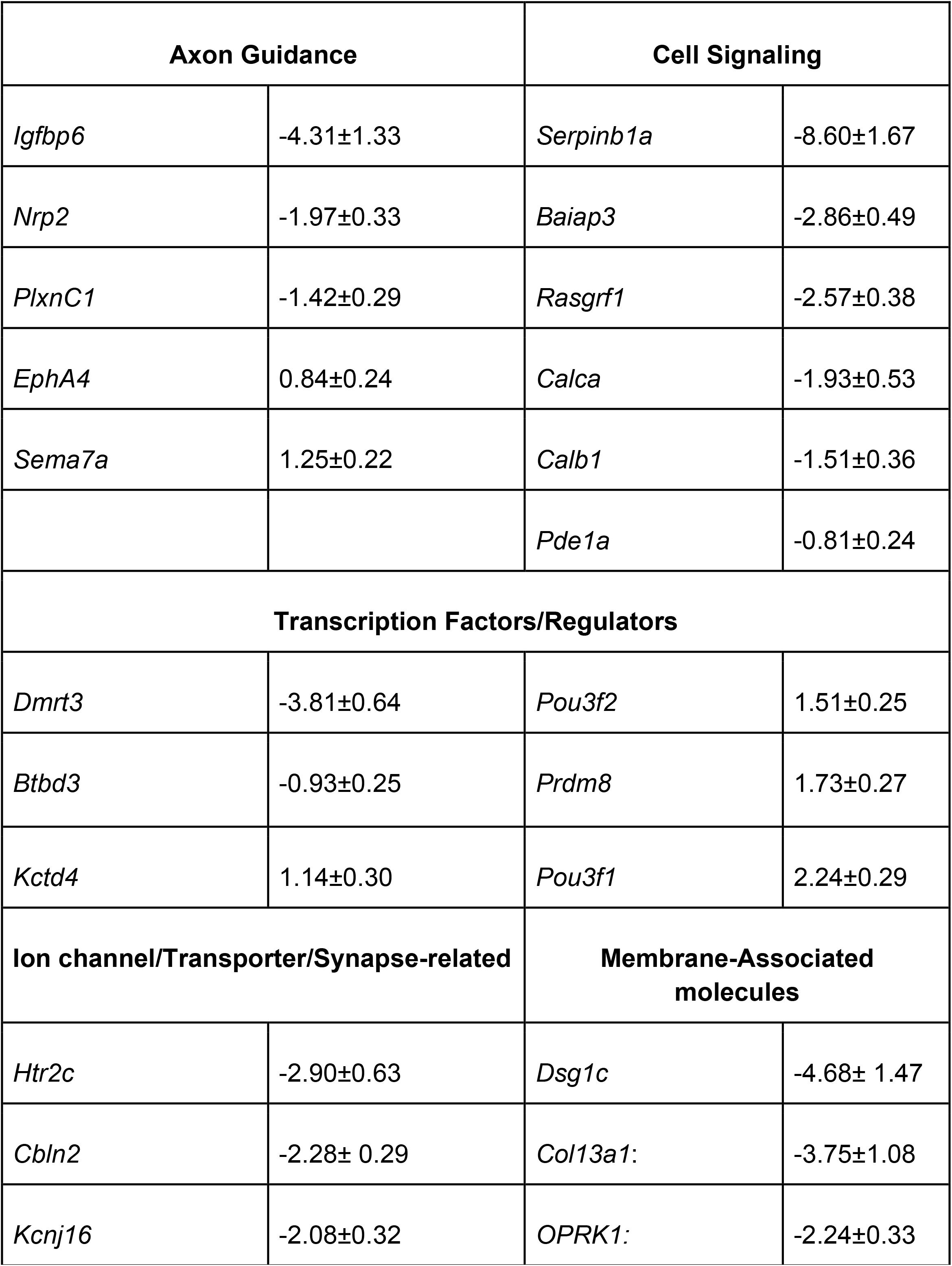
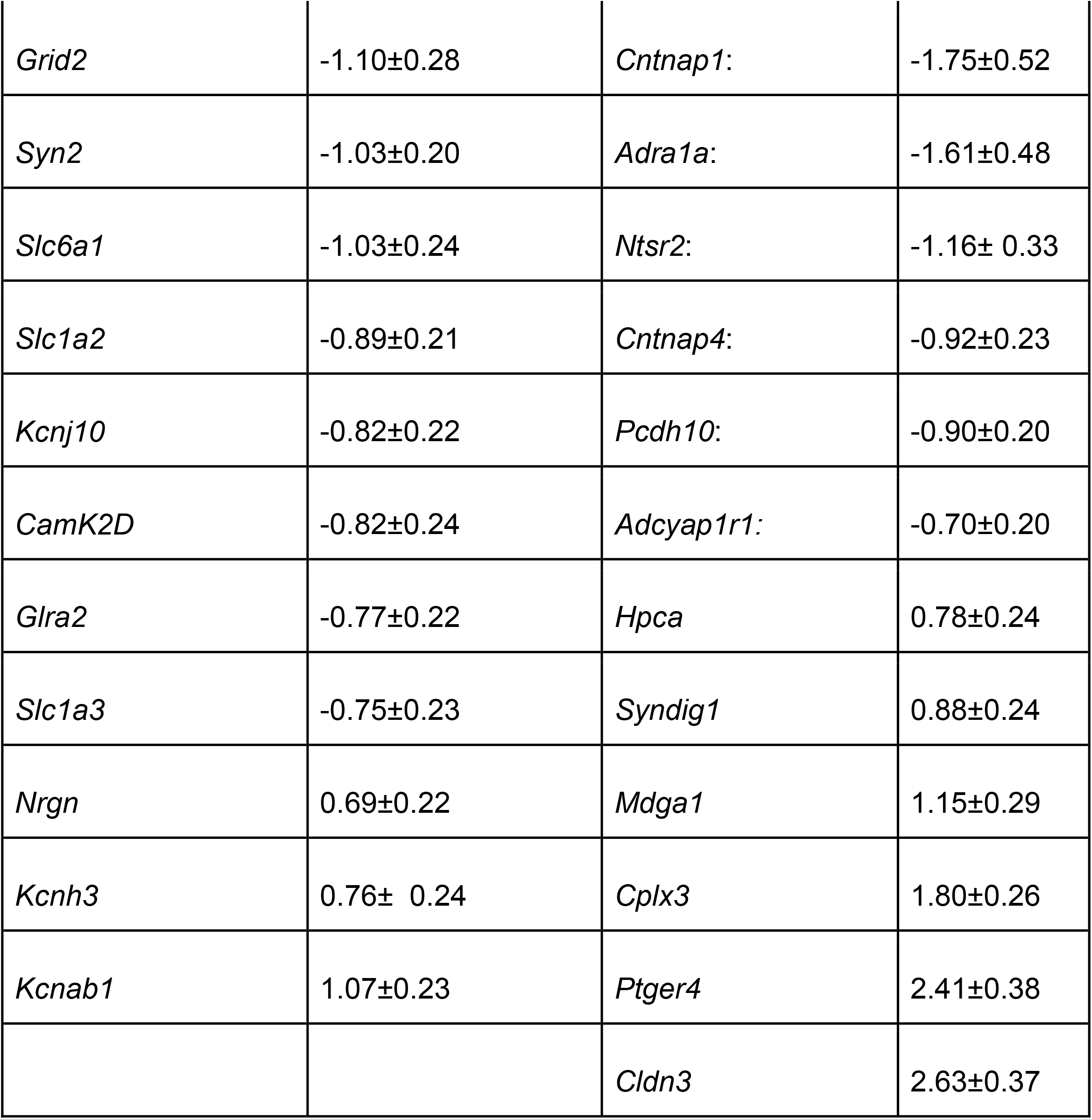
RNA-seq log_2_fold change values (as mean± SEM) of representative differentially expressed genes.

The RNAseq analysis indicated that several genes involved in the bioelectrical properties of neurons were dysregulated in the *Emx1Cre*^*11*.*5*^;*Lhx2*^*lox/lox*^ subplate, e.g. a variety of potassium channel subunits encoded by *Kcn* genes and an ionotropic glutamate receptor encoded by *Grid2*. Subplate neurons have been shown to be electrically active as early as E16, and they form transient glutamatergic synapses on migrating multipolar neurons that will populate the cortical plate (27). These observations motivated an examination of whether the electrophysiological properties of subplate neurons were impaired upon Emx1Cre-driven loss of Lhx2. We prepared live slices from control and *Emx1Cre*^*11*.*5*^;*Lhx2*^*lox/lox*^ littermate embryos at stages at the beginning and the end of the waiting period, E15.5 and P0/P1. Subplate neurons were identified by their location between the cell-dense cortical plate with radially oriented neurons and the cell-sparse white matter, and their typical horizontal bipolar morphologies (28). First, we examined the intrinsic electrical properties of control and mutant subplate cells. Most control E15.5 subplate cells displayed a characteristic spontaneous firing pattern with only 36% being silent. In contrast, a larger fraction of mutant subplate neurons were silent (88%; Figure 7A). The resting membrane potentials (RMPs) of mutant and control subplate neurons were similar at E15.5. While control neurons became hyperpolarized over time, mutant neurons failed to do so; by birth, their RMP was more depolarized compared to controls (Figure 7C, D). The input resistance (IR) was also similar at E15.5. However, only control IR decreased by birth whereas mutant subplate cells continued to display a higher IR (Figure 7E, F). Since depolarized RMP and high IR are features associated with immature neurons (29,30), the *Lhx2* mutant subplate appeared to be electrically immature at birth.

**Figure 7:**
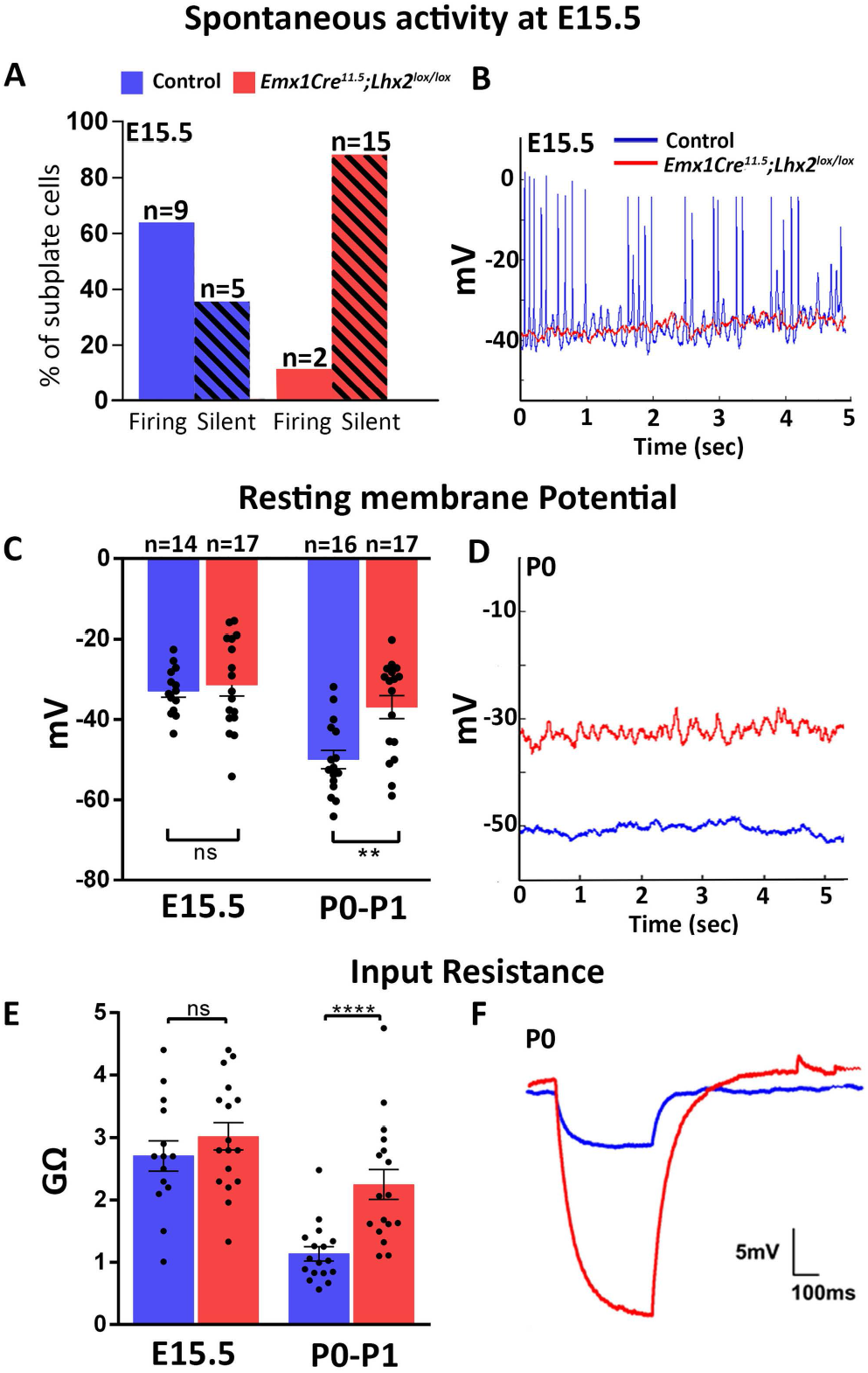
Lhx2 deficient subplate neurons are electrically silent at E15.5 and electrophysiologically immature up to birth. (A-F) Spontaneous responses from slices of E15.5 control and *Emx1Cre*^*11*.*5*^;*Lhx2*^*lox/lox*^ brains. (A) The majority of control neurons fire regularly in absence of any current input, whereas the majority of mutant neurons are electrically silent. (B) Typical examples of mutant and control traces. (C-F) Resting membrane potential (RMP) and Input Resistance (IR) of subplate neurons from E15.5 and P0 control and *Emx1Cre*^*11*.*5*^;*Lhx2*^*lox/lox*^ slices. Both parameters are similar in mutant and control neurons at E15.5 but control neurons display a hyperpolarized RMP and lower IR by birth, which the mutant neurons do not (C, E). For E15.5, n=14 cells were examined from 7 Control brains and n= 17 cells from 10 *Emx1Cre*^*11*.*5*^;*Lhx2*^*lox/lox*^ brains. For P0-P1, n=16 cells were examined from 5 Control brains and n= 17 cells from 5 *Emx1Cre*^*11*.*5*^;*Lhx2*^*lox/lox*^ brains. Data is represented as mean ± SEM. Individual values and statistical analyses are detailed in Table 3. **p≤0.01; ****p≤0.0001. (D, F) Typical examples of mutant and control traces at P0.

Excitability, as assessed by measuring responses evoked by stepwise current injections, was reduced in *Emx1Cre*^*11*.*5*^;*Lhx2*^*lox/lox*^ subplate neurons compared with controls. At E15.5, the mutant subplate neurons were mostly limited to single spikes for input currents from 5-60 pA, whereas wild-type cells displayed a regular spiking pattern (Figure 8A). At P0, though mutant subplate neurons displayed an improved spike frequency, it was lower than that of control neurons at input currents greater than 35 pA (Figure 8B). At both ages, the mutants displayed a higher proportion of single spiking neurons compared to regular spiking neurons (81% vs 21% at E15.5 and 33% vs none at P0; Figure 8C). Mutant subplate neurons also displayed consistently lower spike amplitudes than controls (Figure 8D). The current required to elicit spiking (rheobase) was higher for E15.5 mutant cells (Figure 8E). After recording, biocytin-fillings revealed typical horizontal bipolar morphologies of subplate neurons in mutants as well as controls, at each age (Figure 8F). In summary, the majority of subplate neurons in *Emx1Cre*^*11*.*5*^;*Lhx2*^*lox/lox*^ brains were electrically silent from E15.5, when the thalamocortical afferents first arrive. Their intrinsic bioelectrical properties failed to mature from E15 to P0, and their evoked responses were deficient, remaining typical of immature neurons at birth.

**Table 3:** Electrophysiological properties of Lhx2 deficient subplate neurons.

| Parameter<br>(Mean± SEM) | Age | Control | *Emx1Cre<sup>11.5</sup>;Lhx2<sup>lox</sup>*<br>*/lox* | Statistical Test |
| --- | --- | --- | --- | --- |
| Resting<br>Membrane<br>Potential<br>(in mV) | E15.5 | -32.88 ± 1.55 | -31.34 ± 2.75 | Unpaired two-tailed Student's t test, ns, $p=0.6514$ |
| | P0 | -49.91 ± 2.30 | -36.90 ± 2.90 | Mann-Whitney non-parametric two-tailed test, **<br>$p=0.0017$ |
| Input<br>Resistance<br>(in GΩ) | E15.5 | 2.71 ± 0.24 | 3.02 ± 0.21 | Unpaired two-tailed Student's t test, ns, $p=0.3457$ |
| | P0 | 1.14 ± 0.11 | 2.25 ± 0.24 | Mann-Whitney non-parametric two-tailed test, ****<br>$p<0.0001$ |
| Spike<br>Amplitude<br>(in mV) | E15.5 | 74.49 ± 5.38 | 55.88 ± 2.63 | Unpaired two-tailed Student's t test, **$p=0.0070$ |
| | P0 | 91.55 ± 1.94 | 82.04 ± 2.29 | Unpaired two-tailed Student's t test,<br>** $p=0.0033$ |
| Rheobase<br>(in pA) | E15.5 | 15.36 ± 3.03 | 22.08 ± 3.28 | Mann-Whitney<br>non-parametric<br>two-tailed test,<br>*$p=0.0350$ |
| | P0 | 28.53 ± 2.3 | 23.75 ± 3.31 | Mann-Whitney<br>non-parametric<br>two-tailed test, ns,<br>$p=0.2497$ |

**Figure 8:**
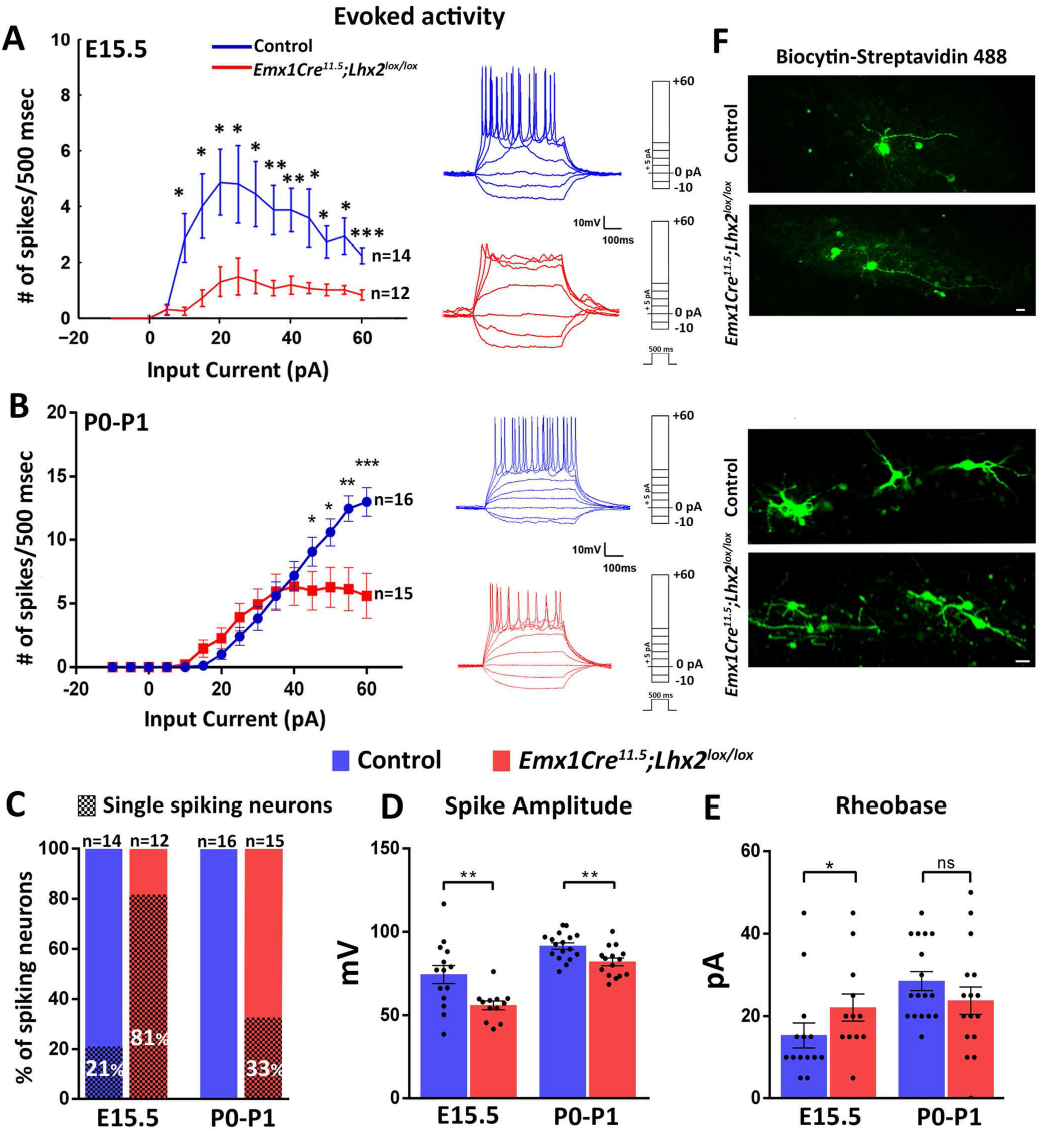
Lhx2 deficient subplate neurons display attenuated evoked responses at E15.5 and birth. (A-E) Evoked responses from slices of E15.5 Control and *Emx1Cre*^*11*.*5*^;*Lhx2*^*lox/lox*^ brains. (A, B) Mutant neurons have an attenuated spike frequency compared to controls at E15.5 and P0. Typical examples of mutant and control traces are shown adjacent to the line graphs. (C) The proportion of single-spiking neurons is greater in mutants compared to controls at E15.5 and birth. (D) Mutant neurons display decreased spike amplitudes at both stages. (E) Current injection required to elicit the first action potential is higher for E15.5 mutant neurons. Data is represented as mean ± SEM. Individual values and statistical analyses are detailed in Table 3. *p≤0.05; **p≤0.01; ***p≤0.001. (F) Biocytin fills of mutant and control subplate neurons at the end of the recording reveals horizontal bipolar morphologies at both ages. Above: E15.5. Below: P0. All scale bars are 10 µm.

In summary, our results revealed dysregulation of diverse set of mechanisms in *Emx1Cre*^*11*.*5*^;*Lhx2*^*lox/lox*^ subplate neurons that could underlie the phenotype of premature exuberant overgrowth by the thalamocortical axons into the cortical plate and an eventual attrition of thalamocortical innervation of the postnatal cortex.

## Discussion

The mechanisms that produce normally functioning subplate neurons are poorly understood. Our study offers insights into this process by highlighting an essential transcriptional regulatory function of Lhx2 that operates in cortical progenitors at E11.5, when the subplate is born.

In order to identify the precise stage of progenitor that requires Lhx2 function, and also to discriminate between progenitors versus postmitotic neurons, we used different Cre lines and compared the resulting phenotype with that seen in *Emx1Cre*^*11*.*5*^;*Lhx2*^*lox/lox*^ brains. NexCre-mediated *Lhx2* recombination does not recapitulate the Emx1Cre-mediated phenotype, since the thalamocortical axons wait at the subplate without showing aberrant innervation and barrels are formed in postnatal stages. When *Lhx2* recombination is induced via CreER, only tamoxifen administration at E11.5, but not E12.5-E13.5, recapitulates the Emx1Cre-mediated phenotype. Therefore, we have uncovered a novel Lhx2 function that selectively operates in E11.5 progenitors and not in postmitotic neurons. Loss of Lhx2 from these progenitors has profound consequences on the transcriptome and the bioelectrical properties of the subplate neurons which are born at E11.5. Since the subplate is the primary target of the thalamocortical pathway when it enters the dorsal telencephalon, we propose that loss of Lhx2 in E11.5 progenitors results in a subplate with deficient molecular properties such that the thalamocortical axons appear to overshoot it and enter the cortical plate prematurely.

### Interaction of thalamocortical and corticothalamic axons in the ventral telencephalon

The subplate plays a guidance role for the entry of the thalamocortical pathway into the dorsal telencephalon and for its initial trajectory below the cortical plate. In the *Tbr1* and *Gli3* mutants, the subplate appears to be absent altogether. The thalamocortical axons extend towards the pallial-subpallial boundary but fail to cross it, suggesting that interactions between subplate projections in the ventral telencephalon are required for thalamocortical afferents to enter the dorsal telencephalon (31–34). Thalamocortical afferents target subplate neurons even when they are mislocalized. In the *reeler* mutant, cortical layering is inverted (35), and the subplate, identified by the expression of CTGF and Cplx3, is located above the cortical plate (a “superplate”). Thalamic axons extend through the cortical plate and reach the superplate and then turn downward to innervate layer 4 (36,37). In the *p35* mutant, subplate neurons are located in the middle of an inverted cortical plate, in which earlier born deep layer neurons are positioned above the subplate and later born superficial layer neurons are positioned below the subplate. The thalamic axons enter the cortex obliquely towards the mispositioned subplate and eventually grow towards the marginal zone (38,39). Together, these studies demonstrate that thalamocortical axons cannot cross the pallial-subpallial boundary without the presence of the subplate and will first target the subplate, even when it is mislocalized, underscoring its importance as a primary target.

In contrast to the mutants described above, in which subplate neurons are either absent or mislocalized, the subplate is correctly positioned below the cortical plate in *Emx1Cre*^*11*.*5*^;*Lhx2*^*lox/lox*^ brains. Corticothalamic projections reach the thalamus similar to wild-type brains, and thalamocortical fibers also navigate normally through the ventral telencephalon and cross the pallial-subpallial boundary. This suggests that the role of the subplate in guiding thalamocortical fibers into the dorsal telencephalon is unaffected. This mutant, therefore, serves as an excellent model to analyze mechanisms that control key subplate properties beyond its guidance function in terms of entry of thalamocortical axons into the dorsal telencephalon.

### VB patterning and cortical area patterning

Previous studies have shown that the VB is responsive to its connectivity with its target cortical area. When the S1 area is shifted to an extreme caudal location upon cortex-specific loss of COUP-TF1, the thalamocortical innervation targets the ectopic S1 and barrels are formed there (40). When the S1 area is reduced as a consequence of cortex-specific loss of *Pax6*, or re-patterned to a motor identity upon cortex-specific loss of Ctip1, the VB undergoes selective re-patterning and/or apoptosis (18, 19). Similar mechanisms could explain why the VB continues to shrink postnatally in the *Emx1Cre; Lhx2*^*lox/lox*^ mutant (Figure 2). Our study extends this target-dependent survival model further, in that VB shrinkage appears to begin embryonically, as a result of the thalamocortical axons encountering a subplate born from Lhx2-deficient progenitors.

### Progenitor-specific mechanisms regulate the properties of the postmitotic subplate

The subplate is described as the most mature neuronal population in the cortex since it is born first and achieves properties associated with mature neurons well ahead of the other cortical neurons [reviewed in (41)]. Synaptic transmission from subplate neurons plays an instructive role in multipolar to bipolar transition of migrating neurons in the developing cortex as early as E16 (27). Thalamocortical axons are also known to make synaptic contacts with the subplate, which serves as a critical and necessary primary target of the thalamocortical pathway (4-5, 9-11, 42, 43). Our study uncovers severe electrophysiological deficits in postmitotic subplate neurons that are born from Lhx2-deficient cortical progenitors from E15.5, when thalamocortical axons contact them. Furthermore, the transcriptome of the subplate arising from *Lhx2*-null progenitors shows gene dysregulation that has little overlap with that seen when Lhx2 is lost postmitotically in those very cells. It is also striking that loss of Lhx2 results in thalamocortical axons displaying inappropriate exuberant overgrowth only in the former but not the latter case. Recent work has shown that transcriptional priming of daughter neurons by their mother cells plays a critical role in setting the developmental trajectories during differentiation (44,45). Our study motivates a broader investigation of transcriptional and epigenetic controls in progenitors that control the properties of the resulting postmitotic progeny, eventually regulating how neurons participate in cortical circuitry.

## Materials and Methods

### Mice

All animal protocols were approved by the Institutional Animal Ethics Committee of TIFR according to regulations devised by the Committee for the Purpose of Control and Supervision of Experiments on Animals (CPCSEA). Noon of the day of vaginal plug was designated as embryonic day 0.5 (E0.5). Mouse embryos of either sex were harvested at E15.5, E17.5, E18.5, P0, P2 and P7. Controls used for each experiment were age-matched littermates. ISH and IHC for each marker, and *in utero* electroporation for each condition, was performed in ≥3 biological replicates. The following lines were obtained from JAX labs: tdTomato reporter Ai9 (strain name: B6.CgGt (ROSA)26Sortm9 (CAG-tdTomato)Hze/J; stock number: 007909), Tamoxifen-inducible CreERT2 line (strain name: B6; 129- Gt (ROSA)26Sortm1 (Cre/ERT)Nat/J; stock number: 004847). The following lines were obtained from: Emx1Cre^11.5^ [Yuqing Li, University of Florida, USA; (46)]; *Lhx2*^*lox/lox*^ [Edwin Monuki, University of California, Irvine, USA; (47)]; NexCre [Klaus Nave, Max Planck Institute for Experimental Medicine; (48)]; TCA-GFP [Takuji Iwasato, National Institute of Genetics, Japan; (17)].

The Emx1Cre^11.5^ line [referred to as Emx1Cre^YL^ in (13)] acts a day later than a commonly used Emx1Cre line that acts from E10.5 [Emx1Cre^10.5^, referred to as Emx1Cre^KJ^ in (13)]. Using Emx1Cre^11.5^ to disrupt *Lhx2* permits the neocortex to be specified (13) instead of being transformed into an ectopic paleocortex, which is seen with Emx1Cre^10.5^ (49). Though reduced in extent compared with controls, the *Emx1Cre*^*11*.*5*^;*Lhx2*^*lox/lox*^ cortex expresses molecular markers of all neocortical layers in the correct order (13). Since Emx1Cre action is specific to the dorsal telencephalon, the Emx1Cre^11.5^ permits the examination of loss of Lhx2 in neocortical progenitors from E11.5 without simultaneously affecting Lhx2 expression in the thalamus, making it an ideal reagent to examine cortex-specific effects on thalamocortical innervation. *Lhx2* is expressed in progenitors and also in newly postmitotic cells [(50); Supplementary Figure S1]. We confirmed that NexCre-mediated recombination of an Ai9 reporter labels the postmitotic subplate (Supplementary Figure S1).

### Sample Preparation

For histological procedures, embryonic brains were harvested in pre-chilled 1X PBS (Phosphate Buffered Saline). Postnatal pups were anesthetized on ice before transcardial perfusion with 4% (wt/vol) paraformaldehyde (PFA; Sigma) made in 0.1 M PB (Phosphate Buffer; pH 7.4). Brains were fixed in 4% PFA overnight at 4°C and equilibrated in 30% sucrose before sectioning.

### Administration of tamoxifen

Tamoxifen (Sigma) dissolved in corn oil (20mg/ml) was administered by oral gavage to pregnant *Lhx2*^*lox/lox*^ dams at different time points as mentioned in the text and embryos were harvested at E15.5. For each embryo, the extent of recombination was examined in one series of sections. Control embryos were littermates with one wild□type copy of the relevant gene. For *Lhx2* ^*lox/lox*^ mice the tamoxifen dosage administered was 75 µg/gm body weight.

### *In situ* hybridization

*In situ* hybridization was performed as follows: the sections were fixed in 4% PFA, washed in 1 X PBS, and treated with Proteinase K (1µg/ml). Hybridization was performed overnight at 70°C in hybridization buffer (4xSSC, 50% formamide, and 10% SDS) containing different anti-sense RNA probes. Post-hybridization washes were performed at 70°C in Solution X (2xSSC, 50% formamide, and 1% SDS). These were followed by washes in 2xSSC, 0.2xSSC, and then Tris-buffered saline–1% Tween 20 (TBST). The sections were incubated in anti-digoxigenin Fab fragments (Roche) at 1:5000 in TBST overnight at 4°C. The color reaction was performed using NBT/BCIP (Roche) in NTMT (100 mM NaCl, 100 mM Tris-pH 9.5, 50 mM MgCl2, and 1% Tween-20) according to the manufacturer’s instructions.

### Probe preparation

All probes were prepared by in vitro transcription using a kit from Roche (Mannheim, Germany) as per manufacturer’s instructions. Templates for *CTGF, Hpca, Lhx2 exon2-3, Nrp2, Nurr1, SERT* were generated by PCR using specific primers from E15.5 (for *Lhx2 exon 2-3, Nurr1*), P0 (*Hpca, Nrp2*) and P7 (for *CTGF, SERT*) mouse brain cDNA (T7 polymerase promoter sequence was added to the reverse primer sequence). Template for *RORβ* was generated from plasmid DNA by restriction enzyme digestion.

CTGF-Fwd: AGCGGTGAGTCCTTCCAAAG

CTGF-Rev: GTAATGGCAGGCACAGGTCT

Hpca-Fwd: CAGGACCTGCGAGAGAACAC

Hpca-Rev: CAGTCCTCTTTTCCGGGGTC

Lhx2 exon2-3-Fwd: CGCGGATCCACCATGCCGTCCATCAGC

Lhx2 exon2-3-Rev: TAATACGACTCACTATAGGG

Nrp2-Fwd: AGAAGCCAGCAAGATCCACC

Nrp2-Rev: GGCCAGACTCCATTCCCAAA

Nurr1-Fwd: CAGTCCGAGGAGATGATGCC

Nurr1-Rev: AACCATCCCAACAGCTAGGC

SERT-Fwd: CAAAACGTCTGGCAAGGTGG

SERT-Rev: CATACGCCCCTCCTGATGTC

### Immunohistochemistry

Brains were sectioned (30 μm) using a freezing microtome and sections were mounted on Superfrost Plus slides (Electron Microscopy Sciences). Sections were quenched with 0.1 M PB (pH 7.4) containing 50 mM NH_4_Cl for 10 mins followed by washing with 0.1 M PB containing 0.1% (vol/vol) Triton X-100 for 10 mins. Sections were incubated in blocking solution containing 10% (v/v) horse serum (Invitrogen) in 0.1M PB with 0.3% (vol/vol) Triton X-100 (Sigma) for 45 min at RT. The sections were then incubated overnight at 4°C in primary antibodies: biotinylated goat anti-GFP (1:400; Abcam, catalog #ab6658), Mouse anti-RFP (1:200; Allele Biotech, catalog #ABP-MAB-RT008, rabbit anti-Cplx3 (1: 200; Synaptic Systems, catalog#122302). The sections were then washed in 0.1 M PB, followed by incubation in secondary antibody for 1.5 hrs at room temperature. Secondary antibodies used were as follows: streptavidin Alexa-488 (1:800; Invitrogen, catalog #S32354) for GFP, Goat anti-mouse antibody conjugated to Alexa-568 (1:400, Molecular Probes, catalog #A11004) for RFP and donkey anti-rabbit antibody conjugated to Alexa-647 (1:400, Molecular Probes, catalog #A31573) for Complexin3. The slides were again washed with 0.1 M PB and mounted in Fluoroshield (Sigma).

### *In utero* electroporation

This procedure was used to visualize the thalamocortical tract at embryonic stages since the TCA-GFP line does not display GFP expression in the thalamocortical axons until postnatal day 2. *In utero* electroporation was performed to transfect a GFP-expressing plasmid into the diencephalon of control and mutant embryos at E11.5, when the neurons of the VB nucleus are born.

All procedures conducted followed the guidelines prescribed by the Institutional Animal Ethics Committee. Timed pregnant *Lhx2*^*lox/lox*^ dams with E11.5 embryos were anesthetized with isoflurane (4% induction, 3% during the surgery) followed by oral administration of Meloxicam (Melonex, United Pharmacies). Uterine horns were successively exposed after a midline laparotomy of 1 cm. Embryos were injected with 1-2 μl plasmid DNA solution dissolved in nuclease-free water with 0.1% Fast Green, into the third ventricle (for targeting the dorsal thalamus) through the uterine wall using a fine-glass microcapillary. pCAGG-IRES-eGFP (2 μg/μl, a gift from G. Fishell, Harvard Medical School) was injected and plasmid DNA was prepared using Macherey Nagel Nucleobond Xtra MaxiPrep Kit (catalog no.740414). Embryos were electroporated by holding their head between tweezers-style circular electrodes (3mm diameter, BTX Harvard Apparatus, catalog #450487) across the uterus wall by delivering three pulses (34 V, 40 ms duration with 999 ms intervals) with a square-wave electroporator (Nepagene CUY21). The uterine horns were returned into the abdominal cavity, the wall and skin were sutured, and the embryos were allowed to continue their normal development until E15.5. The uterine horns and the abdominal cavity were kept moist throughout the surgery (20-25 mins) by flushing 0.9% NaCl (prewarmed to 37°C). Animals were kept on a 37°C warm plate for 30 mins for postsurgical recovery. An oral suspension of Meloxicam (Melonex, United Pharmacies) was mixed with the water in the feeding bottles of the dams (0.6 μl/ml) as an analgesic and given to the animals until 2 days after surgery.

### Imaging

Bright-field images were taken using a Zeiss Axioplan 2 + microscope, Nikon Digital Sight DS-F12 camera, and Nikon NIS 4.0 imaging software. Images of immunohistochemistry slides were obtained using a Zeiss LSM 710 and LSM 880 imaging system. Image stacks were processed using ImageJ (NIH) ((51) and ZEN 2.1 (black) software. Figure panels were prepared using Adobe Photoshop CS6 and Adobe Illustrator.

### Microdissection

P0 brains were dissected in ice-cold HBSS, and 200 µm-thick sections were cut using a vibratome (VT1000S). The lowermost part of the cortical plate (presumptive subplate) was visually identified and micro-dissected under stereomicroscopic guidance in ice-cold HBSS under RNase-free conditions. Microdissected subplate tissue was stored in Buffer RLT Plus (Qiagen) at -80°C and RNA was extracted using RNeasy PLUS Micro kit (Qiagen). Microdissected tissue from 2 brains was pooled to make one replicate sample. Two samples each of control, *Emx1Cre*^*11*.*5*^;*Lhx2*^*lox/lox*^ and *NexCre;Lhx2*^*lox/lox*^ were processed for RNA sequencing after ascertaining that their RIN values were >8.

### RNA sequencing

RNA sequencing was performed by Medgenome Technologies, Bangalore. mRNA was isolated from total RNA, cDNA was prepared, and libraries were prepared using the IlluminaTruSeq RNA Sample preparation V2. Libraries were multiplexed and sequenced as 100 base-pairs (bp) pair-end reads using the Illumina HiSEQ2500 platform, at an expected depth of 60M single reads. Sequenced reads were aligned on the latest mouse reference genome assembly (mm10) using HISAT2, SAM/BAM files were further processed using SAMtools and the number of reads per transcript was calculated and normalized by DESeq2 using R software (52). Genes were considered significantly differentially expressed for FDR of <0.1. For representation, genes with fold change cutoff of +/-1.5 were used. Mutually exclusive and overlapping genes between datasets were identified using Venny (2.1.0). Gene ontology analysis was performed using Gene Ontology Resource (http://geneontology.org/). The RNA-seq data generated in this study will be made available upon publication of the manuscript.

### Electrophysiology

Brain slices were prepared as described previously (25). At least 3 control and mutant brains were used for each age. Briefly, E15.5 embryos and P0-P1 *Lhx2*^*lox/lox*^ and *Emx1Cre*^*11*.*5*^;*Lhx2*^*lox/lox*^ mouse pups were anesthetized by hypothermia and decapitated. Coronal slices (400 µm thick) including the primary somatosensory cortex were cut in ice-cold artificial cerebrospinal fluid [aCSF; (28)] on a vibratome (VT1200S). Slices were allowed to equilibrate in oxygenated aCSF for 30 minutes in a storage chamber before being transferred to the submerged type recording chamber at 30°C-34°C. During preparation and recording procedures, slices were maintained in aCSF containing (in mM) 124 NaCl, 26 NaHCO_3_, 3 KCl, 1.6 CaCl_2_, 1.8 MgCl_2_, 1.25 NaH_2_PO_4_, and 20 D-glucose, pH 7.4, after equilibration with 95% O2 -5% CO2 (osmolarity 333 mOsm). The subplate neurons were visualized using infrared differential interference contrast (DIC) optics using a 40X objective (water immersion lens, 0.9 NA) on an Olympus BX1WI microscope. They were identified by their location, morphology, and electrophysiological properties (25). Subplate neurons had typical horizontal bipolar morphologies and were located between the cell-dense cortical plate with radially oriented neurons and the cell-sparse white matter. Whole-cell patch clamp recordings in current clamp mode were obtained with an Axopatch 200B amplifier (Axon Instruments) and patch-pipettes (3–5 MΩ resistance) filled with (in mM): K-gluconate 155, MgCl_2_ 2, NaHEPES 10, Na-PiCreatine 10, Mg_2_-ATP_2_ and Na_3_-GTP 0.3, pH 7.3 (300 mOsm) and 2 mg/ml Biocytin (Sigma) for post-hoc anatomical staining. Series resistance and input resistance were continuously monitored during the experiment, and the cell was discarded if it changed by >25%. Slices were perfused at all times with aCSF at a flow rate of 2-3ml/min. Electrophysiological Recordings were digitized using Digidata 1400A. Current clamp recordings were filtered at 10 KHz, sampled at 20 KHz and recorded at gain 1. All analysis was done using custom written code in MATLAB (R2013a) and Clampfit (10.7.0.3), and plots were made in GraphPad Prism software.

### Quantification and Statistical Analysis

Statistical tests were performed using Graphpad Prism software version 8.3.0 (538) for Windows, GraphPad Prism Software, San Diego, California, USA. Statistical comparisons were performed after assessing the normality distribution of the data (using Kolmogorov-Smirnov, Shapiro-Wilk and D’Agostino-Pearson omnibus normality tests). Statistical significance was determined using unpaired two tailed Student’s t-test or Mann-Whitney non-parametric test (when data failed normality tests). For samples having unequal variance, Welch’s correction was applied. Multiple comparison analysis was done using a one-way ANOVA test with Holm-Sidak’s multiple comparisons test or Kruskal-Wallis test with Dunn’s multiple comparisons test (when data failed normality tests). All the data was plotted as mean ± SEM. Results were indicated as significant if the p-value is p≤0.05 (*); p≤0.01 (**); p≤0.001 (***); p≤0.0001 (****).

For **Figure 1D**, the fluorescent intensity of axonal terminals marked by TCA-GFP was measured in coronal sections, using ImageJ. Every third (30 μm) section from control and mutant hemispheres, for a total of 3 sections spanning the somatosensory cortex, were examined in each hemisphere. Fluorescence intensity was measured in an area of fixed size (as shown in the cartoon). Statistical test: Unpaired two-tailed Student’s t-test with Welch’s correction, \*\**p*= 0.0034; n=3 for both control and *Emx1Cre*^*11*.*5*^;*Lhx2*^*lox/lox*^ brains.

For **Figure 2C**, the VB area was measured in 3 hemispheres per age from E17.5-P7, in *Emx1Cre*^*11*.*5*^;*Lhx2*^*lox/lox*^ mutant and control. Every third (30 μm) section from control and mutant hemispheres was processed for *in situ* hybridization for *SERT* expression. A total of 3 sections spanning the rostro-caudal extent of the VB were quantitated using ImageJ, and the values summed from each hemisphere. The average mutant value was normalized to that of the average control value. Statistical test: Ordinary one-way ANOVA with Fisher’s LSD test, F= 10.34, \*\*\*\**p*<0.0001; E17.5 Control vs. Mutant, \**p*=0.0326; E18.5 Control vs. Mutant, \**p*=0.0362; P0 Control vs. Mutant, \*\**p*=0.0043; P2 Control vs. Mutant, \*\*\**p*=0.0003; P7 Control vs. Mutant, \*\*\*\**p*<0.0001.

For **Figure 3F**, the cortical thickness was measured using ImageJ (n=3 brains for each condition). The mutant value was normalized to that of the control value. Statistical test: Ordinary one-way ANOVA with Holm-Sidak’s multiple comparisons test, F=0.6267, ns, *p*=0.6007; Control vs. *Emx1Cre*^*11*.*5*^; *Lhx2*^*lox/lox*^, ns, *p*>0.9999; Control vs. *NexCre;Lhx2*^*lox/lox*^, ns, *p*=0.8440; *Emx1Cre*^*11*.*5*^;*Lhx2*^*lox/lox*^ vs. *NexCre;Lhx2*^*lox/lox*^, ns, *p*=0.8440.

For **Figure 3G and 4G**, the area occupied by aberrant axonal branches (pink solid lines in the cartoon) was normalized to the area of the cortical plate (dark gray region in the cartoon) in mutant and control sections, measured using ImageJ (n=3 electroporated brains per condition).

### Statistical tests

Figure 3G: Ordinary one-way ANOVA with Holm-Sidak’s multiple comparisons test, F=67.14, \*\*\*\**p*<0.0001; Control vs. *Emx1Cre*^*11*.*5*^;*Lhx2*^*lox/lox*^, \*\*\*\**p*<0.0001; Control vs. *NexCre;Lhx2*^*lox/lox*^, ns, *p*=0.9992; *Emx1Cre*^*11*.*5*^;*Lhx2*^*lox/lox*^ vs. *NexCre;Lhx2*^*lox/lox*^, \*\*\*\**p*<0.0001. Figure 4G: Ordinary one-way ANOVA with Holm-Sidak’s multiple comparisons test, F=67.14, \*\*\*\**p*<0.0001; Control vs. *Emx1Cre*^*11*.*5*^;*Lhx2*^*lox/lox*^, \*\*\*\**p*<0.0001; Control vs. *CreERT2;Lhx2*^*lox/lox*^ Tam E11.5, \*\*\*\**p*<0.0001; *Emx1Cre*^*11*.*5*^;*Lhx2*^*lox/lox*^ vs. *CreERT2;Lhx2*^*lox/lox*^ Tam E11.5, ns, p=0.9969; *CreERT2;Lhx2*^*lox/lox*^ Tam E11.5 vs. *CreERT2;Lhx2*^*lox/lox*^ Tam E12.5, \*\*\*\**p*<0.0001; *CreERT2;Lhx2*^*lox/lox*^ Tam E11.5 vs. *CreERT2;Lhx2*^*lox/lox*^ Tam E12.75, \*\*\*\**p*<0.0001; *CreERT2;Lhx2*^*lox/lox*^ Tam E11.5 vs. *CreERT2;Lhx2*^*lox/lox*^ Tam E13.5, \*\*\*\**p*<0.0001; *CreERT2;Lhx2*^*lox/lox*^ Tam E12.5 vs. *CreERT2;Lhx2*^*lox/lox*^ Tam E12.75, ns, *p*=0.4386; *CreERT2;Lhx2*^*lox/lox*^ Tam E12.5 vs. *CreERT2;Lhx2*^*lox/lox*^ Tam E13.5, ns, *p*=0.3098; *CreERT2;Lhx2*^*lox/lox*^ Tam E12.75 vs. *CreERT2;Lhx2*^*lox/lox*^ Tam E13. 5, ns, *p*=0.9989).

## Acknowledgments

We gratefully acknowledge the kind gifts of mouse lines from Ed Monuki (*Lhx2*^*lox/lox*^), Yuqing Li (Emx1Cre), Klaus Nave (NexCre). We also thank C. Ragsdale (*RORβ*) for gifts of plasmid DNA used for generating RNA probes and G. Fishell for pCAGG-IRES-eGFP plasmid DNA for *in utero* electroporation; Shital Suryavanshi and the animal house staff of the Tata Institute for Fundamental Research (TIFR) for excellent support; Achira Roy, Anindita Sarkar, Ashwin Shetty, and Hari Padmanabhan for critical input on the manuscript. This work was supported by a Wellcome Trust-Department of Biotechnology India Alliance Early Career Fellowship IA/E/11/1/500402 (GG); a grant from MEXT 16H06459 (TI); a grant from the Department of Biotechnology; Ministry of Science and Technology, India, PR8681 and funds from the Department of Atomic Energy, Government of India, under project number RTI4003 (ST).

---

**Supplementary Figure 1.**
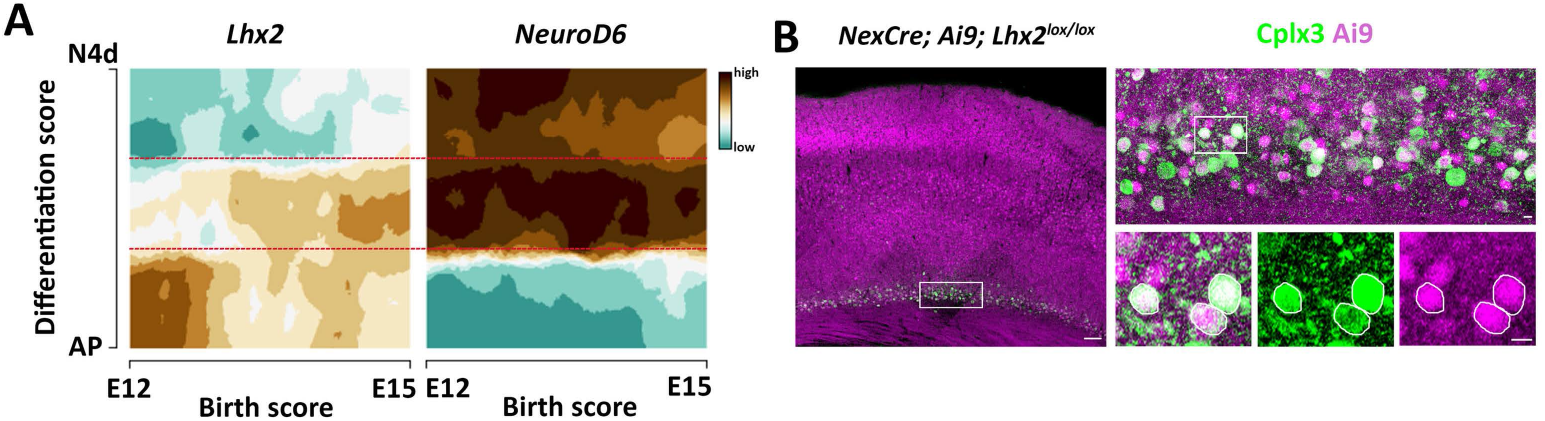
(A) Transcriptional landscape for *Lhx2* and *NeuroD6 (Nex)* in the developing cortical plate of the mouse brain usinghttp://genebrowser.unige.ch/telagirdon/ shows that Lhx2 and NeuroD6 (Nex) are co-expressed in the newly postmitotic population of cortical neurons (region between dashed red lines). (B) An Ai9 reporter recombined using NexCre reveals Ai9 and Cplx3 co-expressing cells in the postmitotic subplate at P7. Scale bars are 100 µm in the low magnification image and 10 µm for the high magnification images.

**Supplementary Figure 2.**
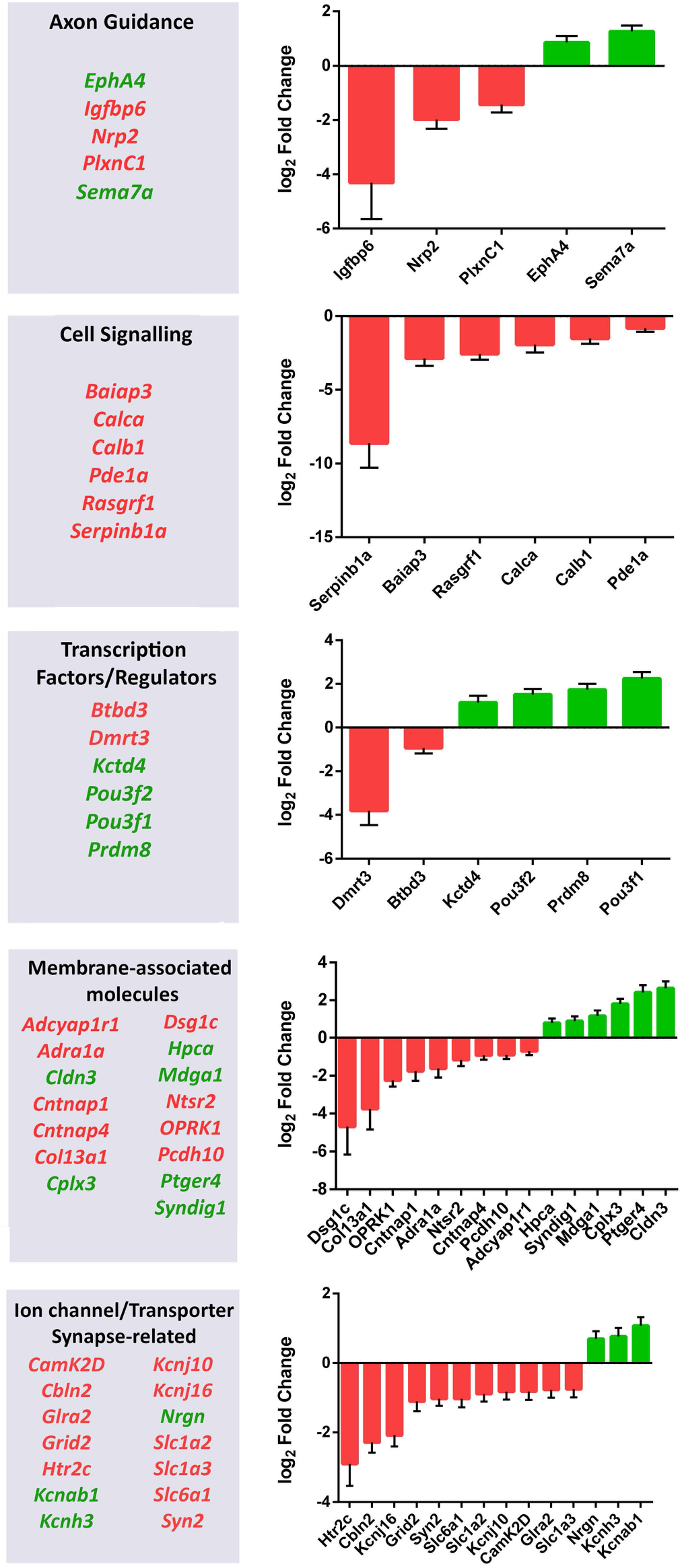
Bar plots representing log_2_fold change values (from RNA-Seq) of selected genes in different functional categories (from Figure 6D).

